# PB-scope: Contrastive learning of dynamic processing body formation reveals undefined mechanisms of approved compounds

**DOI:** 10.1101/2025.06.14.659731

**Authors:** Dexin Shen, Qionghua Zhu, Xiquan Pang, Xian Yang, Dongzhen Pan, Mengyang Zhang, Yanping Li, Zhiyuan Sun, Liang Fang, Wei Chen, Tatsuhisa Tsuboi

**Affiliations:** Institute of Biopharmaceutical and Health Engineering, Tsinghua Shenzhen International Graduate School, Shenzhen 518055, China; Shenzhen Key Laboratory of Gene Regulation and Systems Biology, School of Life Sciences, Southern University of Science and Technology, Shenzhen 518055, China; Department of Systems Biology, School of Life Sciences, Southern University of Science and Technology, Shenzhen 518055, China; Guangming Advanced Research Institute, Southern University of Science and Technology, Shenzhen 518055, China; Tsinghua-SIGS & Jilin Fuyuan Guan Food Group Joint Research Center, Tsinghua Shenzhen International Graduate School, Shenzhen 518055, China

**Keywords:** phenotypic screening, deep learning, drug discovery, P-body, membraneless organelles

## Abstract

Membraneless organelles (MLOs) formed by liquid-liquid phase separation contribute to intracellular compartmentalization of specific biological functions. The dynamic regulation of MLOs is critical for RNA metabolism, stress response, and signal transduction. While the number and size of MLOs serve as valuable indicators of cellular states, leveraging their spatial distribution and diverse quantitative characteristics to infer cellular conditions has posed significant challenges. Here, we present processing body (PB)-scope: an unsupervised deep learning-based framework designed for imaging-based phenotypic screening of PB, one representative MLO. The model was trained on a substantial dataset comprising over 400,000 single-cell confocal microscopy images from a colon cancer cell line treated with 280 compounds. PB-scope enabled precise drug classification based on their effects on PB characteristics, including number and size. This artificial intelligence framework offers insights previously obscured by the dynamic and subtle nature of PBs. Notably, PB-scope identified drugs with a shared function in the JAK signaling pathway, a previously unrecognized regulator of PB dynamics. PB-scope holds the potential for adapting to the study of other MLOs, offering broad applicability in this emerging field of cell biology.

## INTRODUCTION

Phenotypic profiling has long been used in drug screening, primarily for compounds with known targets^1–5^. Recent advances in computational image recognition have enabled the development of novel methodologies for identifying previously unrecognized drug targets^6^. In particular, phenotypic screening strategies that leverage deep learning models applied to large-scale fluorescent imaging data have shown promise in analyzing complex biological features. These models can uncover patterns and correlations that are often difficult to detect using traditional approaches, offering new insights into the mechanisms underlying disease progression and drug responses^7–11^. Recent studies have shown that a cellular morphology-based Convolutional Neural Network (CNN) system can serve as a powerful tool for anti-senescence drug screening^10^. It has also been shown that the nuclear morphology of human fibroblasts can predict senescence with an accuracy of up to 95% by utilizing Deep Neural Network (DNN)^12^. However, these supervised approaches rely on well-characterized cell models or cells treated with drugs of known mechanisms of action (MOAs) as training labels.

In recent years, unsupervised learning has been widely employed in clustering tasks, as it does not rely on labeled data^13–16^. Among the most influential frameworks in this domain is Simple Framework for Contrastive Learning of Visual Representations (SimCLR) ^17^, which revolutionizes self-supervised representation learning by maximizing agreement between augmented views of the same image while minimizing similarity with other instances in the batch. This contrastive learning approach enables the model to extract robust and discriminative features without requiring annotations, allowing for the identification of hidden structures and patterns within the data, making it ideal for exploratory analysis. They have been particularly effective in profiling phenotypic features of extensive cellular structure, such as mitochondria ^18^. As organelles central to cellular energy production, mitochondria exhibit intricate network structures in microscopic images, and changes in their morphology have been linked to various diseases and cellular stress responses. Such phenotypic changes provide a rich source of information for drug screening^19–21^. Recent studies have demonstrated the potential of self-supervised models that utilize re-identification networks to identify MOAs by capturing mitochondrial phenotypic alterations^22^. Despite these promising developments, it remains uncertain whether phenotypic drug screening can be extended to membraneless organelles (MLOs), given their highly dynamic assembly and disassembly.

MLOs are dynamic subcellular compartments in eukaryotic cells formed through liquid-liquid phase separation of proteins and/or RNAs from the surrounding milieu^23,24^. Recent studies highlight their significance in diverse pathophysiological conditions^25,26^. Cytoplasmic processing body (P-body), as one representative MLO, is evolutionarily conserved from yeast to humans. It consists of translationally repressed mRNAs and numerous proteins associated with mRNA deadenylation and decapping, as well as mRNA translation repression ^27,28^. P-bodies are highly dynamic, with posttranslational modifications of their core proteins, including phosphorylation and ubiquitination, influencing their turnover^29–31^. However, the mechanistic details of their assembly and disassembly remain largely elusive. While prior studies have focused on changes in P-body size and number, other features, such as subcellular positioning, may provide additional insights but pose technical challenges for accurate quantification.

To address this, we developed PB-scope, an image-based screening platform that captures comprehensive P-body features at single-cell resolution, enabling the identification of small molecules that influence P-body assembly/disassembly. This unsupervised deep-learning framework employs contrastive clustering to analyze over 400,000 segmented single-cell images acquired using a spinning disk confocal microscope (200 nm resolution) from a colorectal cancer cell line treated with 280 compounds. Using in-house segmentation models, we validated that the PB-scope effectively integrates multiple P-body-related features, facilitating accurate compound grouping. This framework leverages the power of artificial intelligence to analyze and interpret complex biological data, offering insights that were previously difficult to obtain due to the dynamic and nuanced nature of MLOs. Notably, our analysis uncovered a class of drugs with a shared MOA, inhibitors of the JAK signaling pathway, previously unknown in P-body regulation. Beyond identifying compounds that regulate PB assembly, PB-scope holds the potential for adapting to the study of other MLOs, offering broad applicability in this emerging field of cell biology.

## Results

### PB-scope: an unsupervised deep learning framework for large-scale phenotypic screening of P-body regulators

The framework of the PB-scope is illustrated in Figure 1. To investigate the *in vivo* dynamics of endogenous P-bodies, we generated a DDX6-GFP knock-in HCT116 cell line for fluorescent labeling of P-bodies. These cells were then treated with 278 FDA-approved kinase inhibitors and two control compounds previously reported to affect P-body assembly, MG132 and thapsigargin (Supplementary Table. 1). Time-lapse high-resolution image data were collected using the CQ1 Benchtop High-Content Analysis System (Yokogawa) at one-hour intervals over eight hours post-treatment (Fig. 1a). We simultaneously monitored P-bodies, mitochondria, nuclei and cell membrane in living cells (Fig. 1b). The mitochondrial channel was used for cell segmentation via Cellpose^32,33^, while the remaining three channels contributed to the construction of a high-quality dataset comprising over 400,000 single-cell images (Fig. 1c). Subsequently, we trained a contrastive clustering model to extract features and cluster phenotypes associated with various drug treatments (Fig. 1d).

**Figure 1.**
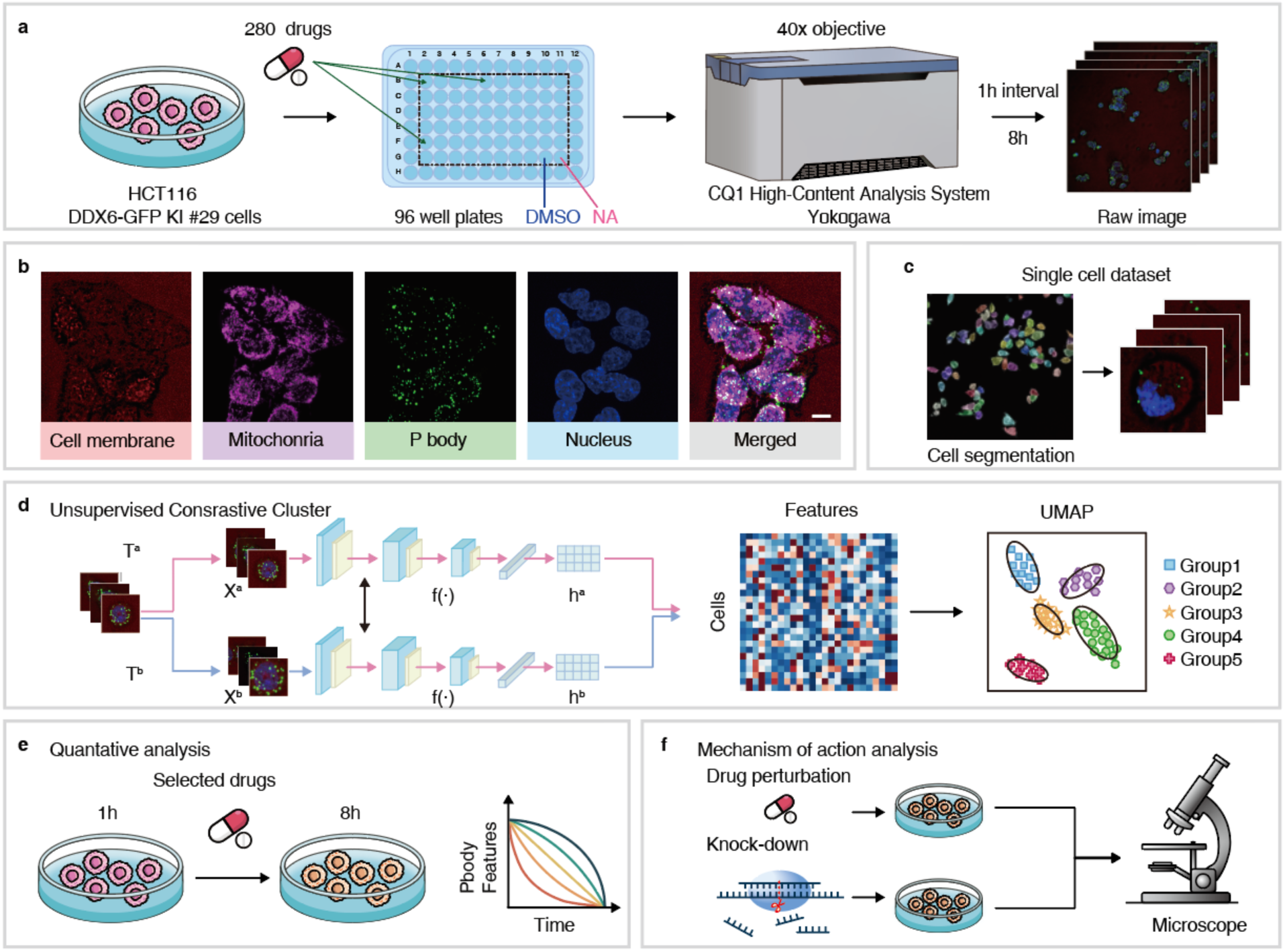
Overview of PB-scope: an unsupervised deep learning-based framework for large-scale phenotypic screening on P-bodies. (a) HCT116 cells stably expressing DDX6-GFP were plated in 96-well plates, treated with 280 compounds at 10µM concentrations, and subjected to high-content imaging using the CQ1 confocal quantitative imaging system. (b) The analyzed images consist of four channels: (i) Bright-field image for cellular morphology, (ii) Mitochondrial network, (iii) Processing body, and (iv) Nucleus. Merged composite demonstrates spatial relationships between these subcellular compartments. Scale bar: 10μm. (c) Mitochondrial channels were processed through Cellpose 3.0 to generate a curated dataset containing over 400,000 high-quality single-cell images. (d) A contrastive clustering framework was implemented for unsupervised feature extraction, followed by UMAP dimensionality reduction to identify compounds with analogous mechanism-of-action (MOA) profiles through cluster localization analysis. (e) Quantitative analysis of P-body formation followed by drug treatment. (f) Mechanistic evaluation of lead compounds via imaging analysis.

PB-scope employs Contrastive Clustering^15^ as the tool for feature extraction (Supplementary Fig. 1). Contrastive Clustering adopts a two-branch architecture that performs data augmentation separately and then utilizes the contrastive learning process to enhance feature representation and clustering accuracy. The model decomposes the clustering task into two distinct yet interconnected levels: cluster-level contrastive learning and instance-level learning. Each row of the feature matrix can be interpreted as a soft label for an instance, representing the probability distribution of the instance belonging to each cluster. Conversely, each matrix column embodies the cluster representation, illustrating how different instances are distributed across clusters and effectively indicating which instances should be grouped together within each cluster. This dual-level approach allows the model to simultaneously refine the representation of data points and their cluster assignments. As a result, the model achieves a more comprehensive understanding of the data, facilitating more accurate and meaningful clustering outcomes.

The choice of data augmentation is critical in contrastive learning, as it significantly influences model performance^17^. In this study, we applied a suite of augmentation techniques optimized for fluorescent microscope data, including standard transformations such as cropping and resizing, horizontal flipping, grayscale conversion, and standard Color Jittering. In addition, we implemented a color space transformation^34^ called Multi-Channel Color jitter (MCjitter) that independently takes the intensity of each fluorescent channel into our augmentation strategy. MCjitter works by independently augmenting the intensity of each fluorescent channel, thereby mitigating the effects of channel-specific noise and preserving the integrity of the underlying biological signals. This augmentation technique proved instrumental in enhancing the model’s ability to learn robust and discriminative features from the fluorescent images. Furthermore, to tackle the data imbalance problem, we introduced focal contrastive loss^35^ into the PB-scope. This loss function is designed to dynamically adjust the weights assigned to different samples based on their difficulty, giving more emphasis to hard-to-classify instances and minority classes. By incorporating MCjitter and the focal contrastive loss function, our contrastive clustering framework achieves a notable accuracy of 63.27%, outperforming baseline comparisons across key evaluation metrics. Specifically, it demonstrated superior performance in Adjusted Rand Index (ARI) and Normalized Mutual Information (NMI), underscoring its effectiveness in capturing meaningful cluster structures and aligning with cluster labels (Supplementary Table 2).

Phenotypic features extracted by the model were projected using UMAP^36^, allowing for the visualization of compound clustering based on the shared MOAs and functional effects on P-body. Our approach was further validated through rigorous quantitative evaluations (Fig. 1e, f). Beyond P-body regulation, this approach establishes a generalizable platform for investigating drug actions through unsupervised phenotypic analysis of MLOs.

### Identify drugs inducing P-body phenotypes via PB-scope

Using our contrastive clustering model, we analyzed 282 conditions (280 drug treatments and DMSO/NA controls) following a 6-hour treatment regimen. For each drug, 30 cell images were randomly selected from the test set to generate UMAP embeddings (Supplementary Fig. 2). From this initial cohort, 42 drugs exhibited distinct spatial clustering in UMAP projection, which were subsequently categorized into five groups based on their distribution in the UMAP embedding space (Fig. 2a). Two independent replicate experiments validated the reproducibility of our approach: five drug groups displayed consistent distribution patterns across both replicate experiments (Fig. 2b-g). Notably, structural analogs, such as T0374/T0374L in Group 1 and T1791/T1791L in Group 5, exhibited spatial proximity in UMAP (Supplementary Fig. 3), aligning well with our expectations that structurally similar compounds often induce the same phenotypic change.

**Figure 2.**
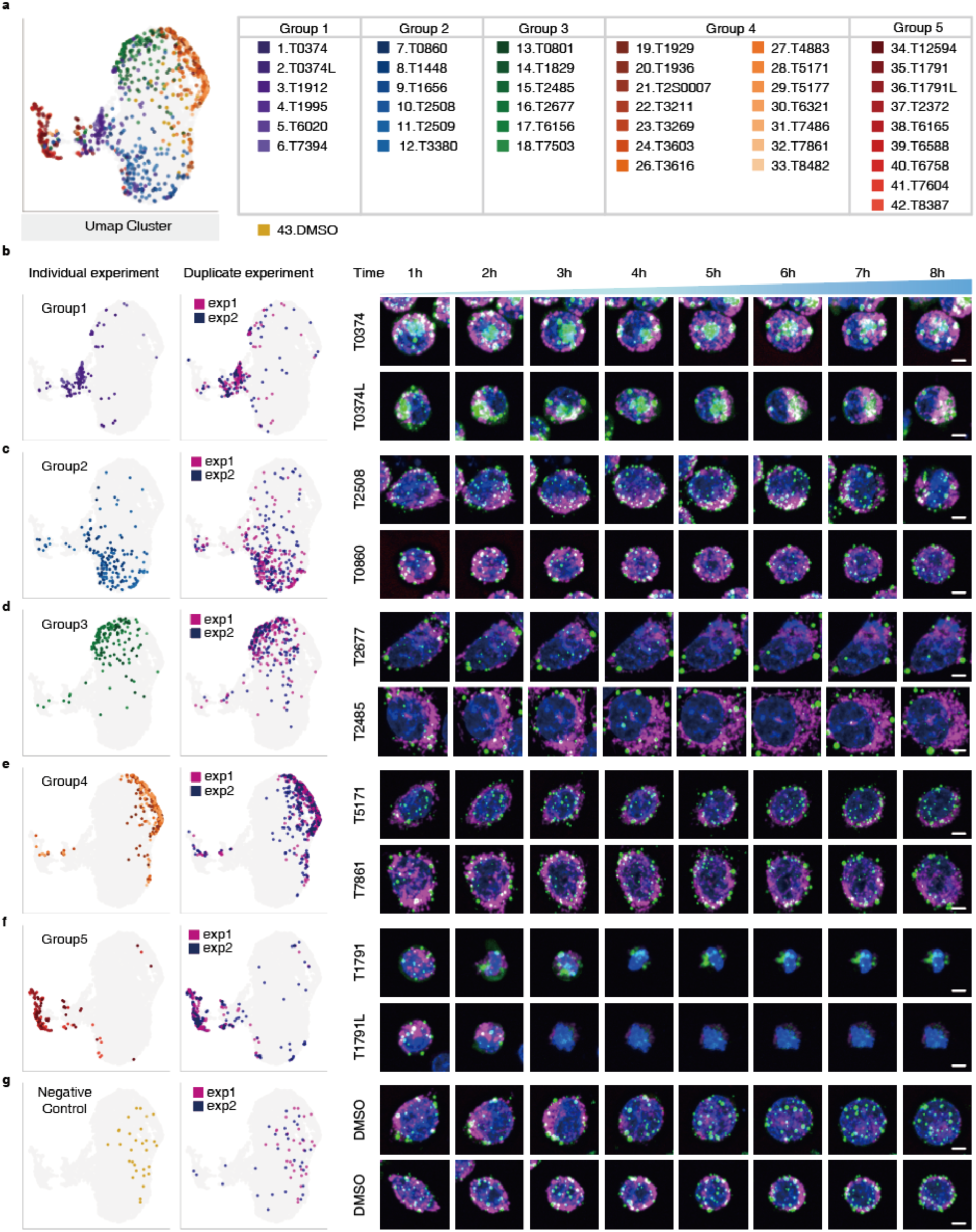
Identification of drugs with analogous P-body phenotypes via PB-scope. (a) UMAP cluster analysis of 42 compounds demonstrates a distinct spatial distribution (negative control: DMSO), based on which five drug clusters were identified. (b)-(f) Experimental validation across two independent biological replicates reveals consistent cluster distribution patterns. Time-lapse montages (0-8 hours, 1-hour intervals) display characteristic phenotypic evolution for each compound group. Scale bar: 5μm. (g) The negative control group exhibited a uniform distribution in the UMAP, which serves as a baseline for comparing the distribution patterns of other assembled groups.

Given the distinctive clustering observed at the 6-hour time point, we further analyzed the data after shorter treatment regimens to see the sensitivity of our method. As shown in Supplementary Fig. 4, PB-scope was capable of distinguishing treatment groups as early as three hours post-treatment, with clustering resolution progressively improving over time. As for the influence on P-body dynamics, we observed that drugs in Groups 1 and 3 led to a reduction in the number of P-bodies, accompanied by an increase in their size (Fig. 2 b, d). In contrast, Groups 2 and 4 showed no visible changes in P-body size or numbers compared to the DMSO control (Fig. 2c, e, g). Additionally, Group 5 induced cellular death around three hours post-treatment (Fig. 2f). These results demonstrate PB-scope’s ability to resolve the full spectrum of drug responses - from subtle P-body modulation to acute cytotoxicity.

### Quantitative analysis of P-body phenotypes

To further investigate the differences among the five drug groups identified through clustering, particularly to explore the latent phenotypic landscape associated with P-bodies in Groups 2 and 4, we performed a quantitative analysis of P-body number and the DDX6-GFP abundance. Given the inherent challenges in manually annotating P-body boundaries with high accuracy, we adopted a simulation-based and deep learning-assisted detection approach to approximate P-body counts and fluorescence intensity (see methods for details, Supplementally Fig. 5, 6). We first generated a synthetic dataset by simulating cells and phase-separated particles (Fig. 3a, Supplementally Figure 5). Cells were modeled as ellipsoids of varying sizes, with background noise following a Gaussian distribution. This simulation dataset was then used to train a YOLOv7 ^37^ network for particle detection (Fig. 3b, c). Before detection, the experimental images were converted to 8-bit format, and their intensity values were normalized to the range of 450 - 1000 using Fiji ^38^. Using this pipeline, we quantified the average number of P-bodies and fluorescent intensity of the DDX6-GFP across compound groups (Fig. 3d-g). (Figure 3d and e are examples from each group. The detailed results are summarized in Supplementary Table 3.) We observed a distinct effect in P-body formation for MG132 and thapsigargin, similar to previously reported: MG132 continuously decreases the number over 8 hours ^41^, while thapsigargin temporarily reduces the number of P-bodies within 6 hours ^39,40^ (Supplementary Fig. 7).

**Figure 3.**
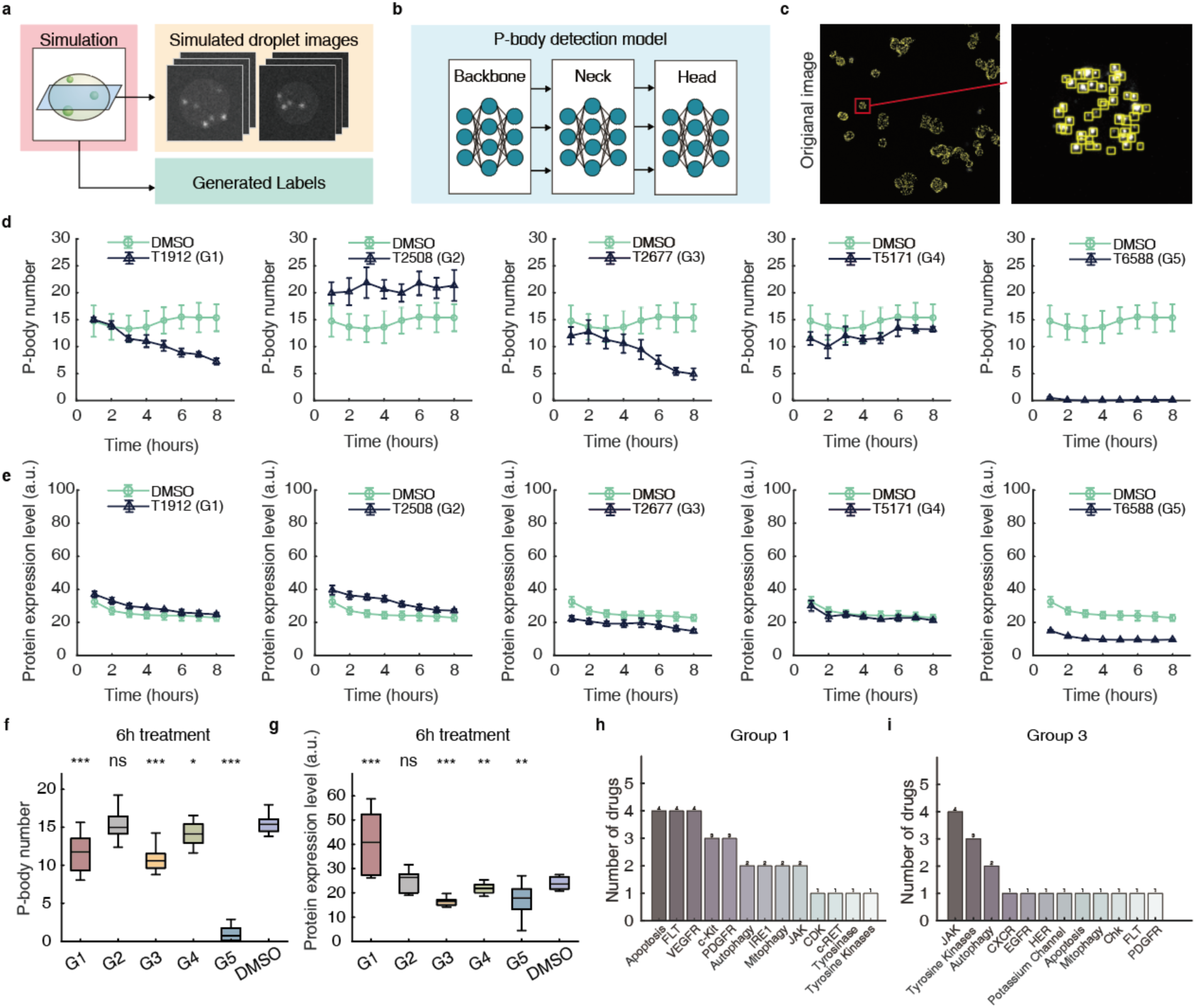
Quantification and mechanisms of action analysis of selected drugs. (a) A simulation model of intracellular p-body was constructed to generate synthetic p-body distributions with ground truth annotations. (b) A YOLO-v7 architecture trained on synthetic datasets was implemented for automated identification and quantitative analysis of P-body formation. (c) Example of P-body detection, achieving >95% agreement with manual annotations (Supplementary Fig. 6). (d) P-body numbers per cell in the time course under different drug treatment groups. (e) DDX6-GFP expression level (a.u.) per cell under different drug treatment groups. Error bars represent the STD of three independent analyses for d and e. (f) Quantitative analysis of P-body numbers at 6 hours post-treatment across different drug groups. (g) Quantitative analysis of DDX6 expression level (a.u.) at 6 hours post-treatment across different drug groups. The *p*-values were determined using the two-tailed Mann–Whitney test for **f** and **g.** The statistical significance compared with DMSO was indicated as *** P < 0.001; ** P < 0.01; * P < 0.05; ns, no significant difference. Data points that lay outside the 15% - 85% range were deemed outliers and excluded from the statistical analysis. (h, i) Mechanism of Action (MOA) profiling for drugs in Groups 1 and 3.

The quantification of P-body numbers reveals a particularly pronounced decline in Groups 1, 3, and 5, with Group 5 exhibiting the most significant decrease in 6-hour post-treatment (Fig. 3f). In contrast to other groups, the quantitative analysis of DDX6 fluorescent intensity shows that Group 1 exhibited a notable increase in fluorescence intensity (Fig. 3g). This trend may be due to the activation of specific intracellular signaling pathways, which promote the fusion of small P-bodies and/or an increase in DDX6-GFP expression, leading to a decline in P-body numbers and an overall enhancement of fluorescence intensity. Conversely, Groups 3 and 5 displayed a significant decrease in cellular fluorescence intensity, albeit through distinct mechanisms. For Group 3, it is likely attributable to downregulated DDX6-GFP expression, while for Group 5, it is driven by drug-induced cell death (Fig. 2f, 3f, g). These analyses also revealed nuanced effects of specific compounds on P-body dynamics, particularly in drug groups that did not exhibit substantial morphological changes under visual inspection (Groups 2 and 4). While Group 2 does not show changes in the number of P-bodies, the variation in P-body number is much higher than compared to the other groups (Fig. 3f). This may be caused by the heterogeneous P-bodies formation, which is difficult to quantify using traditional methodologies. Additionally, Group 4 exhibits a slight decrease in the number of P-bodies. These findings highlight PB-scope’s unique sensitivity in identifying compounds that induce nuanced phenotypic changes, which would be overlooked in standard screening paradigms.

To see whether the observed phenotype fits with the known MOA of the drugs, we checked the known targets of different compounds groups (Fig. 3h, i, Supplementary Fig. 8). Interestingly, apoptosis-related pathways such as FLT and VEGFR were frequently identified as targets in Group 1 (Fig. 3h). Indeed, Group 5 exhibited an apparent apoptotic phenotype, and Groups 1 displayed close geometric proximity to Group 5 in the UMAP embedding (Fig. 2). Furthermore, JAK inhibitors stand out in both Group 1 and 3, which showed opposite effects in intensity, but similar trend on number (Fig. 3h, i). Together, these observations underscore the biological coherence and robustness of our clustering framework. These findings validate that the drugs identified through clustering methods indeed induce changes in cellular phenotypes, thereby confirming the reliability of the screening approach. While visually observed changes in cellular phenotypes provide an intuitive reflection of the overall physiological state of cells under drug influence, quantitative analysis could gain a more detailed profile of even nuanced phenotypic changes.

### The JAK signaling pathway is involved in P-body regulation

To further understand the influence of inhibitors on P-body, we focused on inhibitors targeting Janus kinase (JAK) as they show different UMAP embedding among Groups 1 and 3 (Fig. 3h, i). When we explored the functional targets of the drugs in Groups 1 and 3, we found that 2 out of 6 and 4 out of 6 inhibit JAK1 and JAK2, respectively. To validate the results, we further conducted immunofluorescence experiments on cells treated with siRNA targeting either JAK1 or JAK2 to clarify the effects of this signaling cascade on the quantitative characteristics of P-bodies (Fig. 4a, b, Supplementary Fig. 9). We observed an increased number of P-bodies in HCT116 cells when JAK1 or JAK2 was knocked down. These observations suggest a potential association between JAK signaling and the inhibition of P-body formation. As the JAK pathway is a general signal transduction pathway related to stress response and transcription (Fig. 4c), the protein composition or overall structural integrity of P-bodies must be altered in the absence of functional JAK1 and JAK2. Thus, our screen revealed a previously unrecognized regulatory role for the JAK signaling pathway in maintaining and regulating P-body assembly (Fig. 4c).

**Figure 4.**
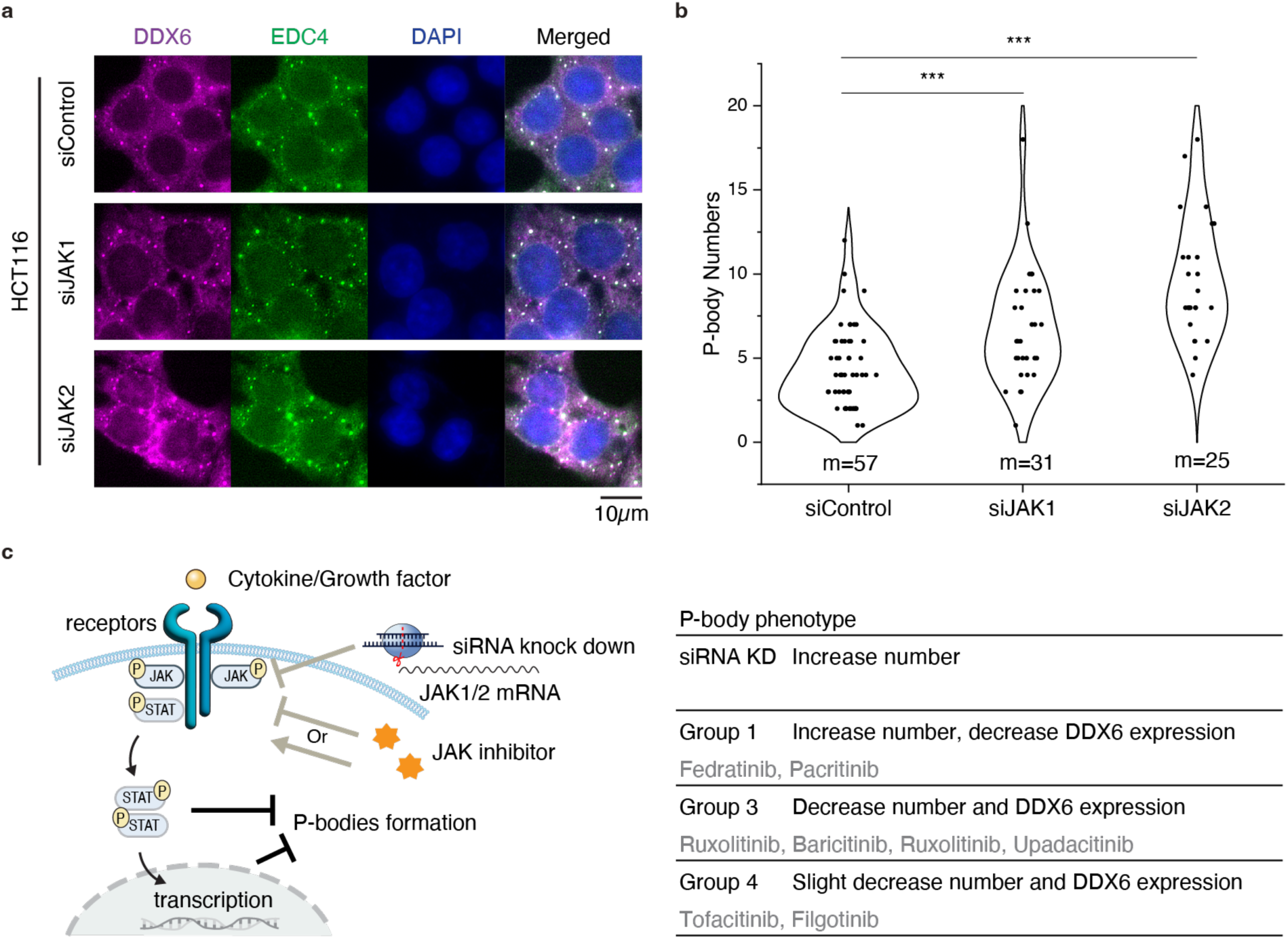
Perturbation of JAK leads to enhanced P-bodies. (a) HCT116 cells were knocked down using JAK1 and JAK2 siRNA, and immunostained for P-body components DDX6 (magenta) and EDC4 (green). The nuclei were visualized with DAPI (blue). Scale bar, 10μm. (b) Quantification of P-body number per cell across three experimental groups. Statistical significance determined by an unpaired t-test was indicated as *** P < 0.001. (c) Model of JAK/STAT signaling pathway-mediated P-body regulation. JAK is activated when cytokines or growth factors bind to their respective receptors, leading to receptor dimerization, JAK and STAT phosphorylation, and subsequent transcriptional regulation. Inhibition of the pathway by knockdown of JAK1/2 leads induction of P-body formation. The JAK inhibitors identified in this work that modulate P-body formation are shown in the right panel.

## Discussion

In this work, we established a robust computational framework for large-scale drug screening based on P-body phenotypes through unsupervised deep learning. We processed and analyzed a dataset comprising over 400,000 microscopic images from HCT116 cells treated with 280 compounds (Fig. 1). The PB-scope is built based on a contrastive clustering framework with two distinct augmentation branches and MCjitter model that provides better UMAP clustering with similar mechanisms of action and pathways (Supplementary Fig. 1 and Fig. 2), enhancing the potential in identifying promising drug candidates. Additionally, we developed a deep learning-based detection model for quantitative analysis of P-bodies and used it to confirm the efficacy and robustness of our initial screening results (Fig. 3). Finally, the identification of JAK pathways in P-bodies regulation was validated with independent perturbation assays (Fig. 4).

Compared to existing phenotypic screening methods^42–44^, the PB-scope presents numerous benefits. First, our approach introduces deep learning for feature extraction and clustering of large-scale image data, enabling unbiased discovery of novel cellular functions. Second, it overcomes the limitations of supervised methods by eliminating the need for labeled data. Moreover, the self-supervised contrastive learning architecture circumvents the constraints of traditional supervised approaches, allows the discovery of compounds with previously uncharacterized mechanisms of action. These technical advances allowed the identification of novel compound clusters showing unexpected P-body modulation effects that warrant further investigation (Fig. 2).

A particularly striking finding was the association between JAK inhibitors and P-body regulation. (Fig. 3h, i, Fig. 4) Our analysis identified JAK1/2 as recurrent targets in Groups 1 and 3, and subsequent validation via siRNA knockdown and immunofluorescence confirmed that JAK inhibition enhances P-body formation. Interestingly, the inhibitors classified in Groups 1 and 3 exhibited distinct P-body phenotypes, such as the induction of larger P-bodies and a reduction in P-body formation, respectively, despite both being JAK inhibitors. These suggest that JAK inhibitors lead to significantly different cellular physiological consequences and require attention to the molecular mechanism of action upon use. Notably, prior studies have demonstrated that P-bodies can be affected by oxidative stress^45^ and viral infection^46^, and the JAK-STAT pathway is one of the core pathways involved in these stress responses. These observations suggest a previously unrecognized functional link between JAK-STAT signaling and P-body dynamics, providing new insights into how kinase pathways influence P-body formation.

While the current framework demonstrates significant advantages over existing methods, three technical constraints require future improvements. First, unsupervised drug screening cannot ensure consistent drug effects across all cells due to intercellular heterogeneity. Future research may need to incorporate more refined cell sorting techniques or develop algorithms that adjust for cellular heterogeneity. Second, unsupervised drug screening lacks precise quantitative criteria. P-bodies show relatively simple morphological features that can be described by size and number. As with mitochondrial phenotype, it is not a typical case that the results of contrastive clustering can be quantifiable. New quantitative methods need to be explored to evaluate the impact of drugs on cells more accurately. Additionally, current methods often require z-projection processing of images, which may result in the loss of valuable information about cell structure and spatial distribution. By utilizing point cloud models such as Pointnet^47,48^ to capture the phenotypic features of cellular structures such as P-bodies and mitochondria, we could preserve the three-dimensional spatial structural characteristics while minimizing the consumption of computational resources.

In summary, PB-Scope has constructed a comprehensive conceptual framework that combines advanced imaging techniques, cell segmentation algorithms, unsupervised comparative learning, and in vitro biochemical experiments. This framework could be easily adapted to analyze other MLOs for phenotypic screening, which could accelerate the process of new drug discovery and development by capturing complex stress responses to various compounds. Future work could expand the dataset and refine deep learning models to enhance accuracy and robustness. Ultimately, our framework has the potential to revolutionize the drug screening process and accelerate the discovery of new and effective therapies.

## Methods

### Construction of cell lines and cell culture

Knock-in (KI) cells were generated from wild-type HCT116 cells. A single-guideRNA (sgRNA) (5’-ttatcaatgttgctcggaat-3’) with overhangs for BbsI (NEB, R3539) restriction was inserted into the pSpCas9(BB)-T2A-Puro (Addgene, 48139). The homologous repair donor template was inserted into the linearized Tet-On 3G vector backbone (courtesy of Dr. Ruijun Tian, Southern University of Science and Technology) using Gibson assembly (YEASEN, 10922ES50). The transfection was carried out using the bicistronic nuclease plasmid with the corresponding donor plasmid at the ratio of 2 to 1, and a total of 9 μg plasmid were transfected in HCT116 cells as described above. The fluorescent-positive single cells were sorted into 96-well plates by FACS to expand single-cell clones. The single-cell clones of homozygous KI cells were selected in which DDX6-GFP replaced WT DDX6. Primers used to genotype single clones are as following: aggtggtatgttctgtgactgt (forword), aggactctgaatttaagtactgcta (reverse).

HCT116 DDX6-GFP KI cells were maintained in Dulbecco’s Modified Eagle Medium (DMEM; Gibco, #10566016), supplemented with 10% fetal bovine serum (FBS; GenClone, #25-550) and 1% penicillin/streptomycin (P/S; Gibco, #15140122), in a humidified atmosphere containing 5% CO2 at 37°C. Prior to passaging the cells onto imaging plates, they were gently washed with DPBS (Gibco, #14190144) to remove any residual medium and detached using 0.05% Trypsin-EDTA (Gibco, #25300054) to facilitate cell dissociation. A total of 12,000 cells per well were seeded into 96-well plates in 200 µL of medium.

### Drug treatments

The library of 278 FDA-approved kinase inhibitor compounds and small molecular compounds MG132 and thapsigargin were purchased from TargetMol (Supplementary Table 1). Cells were cultured in 96-well plates for 48 hours before drug treatment. All wells were treated with a 10μM concentration of the drugs. The live cell imaging data were collected over 8 hours. Dimethyl sulfoxide (DMSO) was set as a negative control to assess the cell phenotype at the basal level.

### Imaging

Live-cell imaging was performed using the CellVoyager™ CQ1 Benchtop High-Content Analysis System (Yokogawa) equipped with a 40× objective lens. Four fields of view were captured per well. Z-stack imaging was conducted with 0.5 µm step sizes across a total depth of 10 µm, utilizing an autofocus routine on the 405 nm channel with 2 µm steps over a 10 µm range. For fluorescent staining, nuclei were labeled with 5 µg/mL Hoechst 33342 (Beyotime, #C1022), and mitochondria were labeled with 50 nM MitoTracker Deep Red FM (Yeasen, #40743ES50). After a 30-minute incubation with the dyes, cells were washed twice with PBS, and fresh medium (Gibco, #21063029) supplemented with 10% FBS (GenClone, #25-550) and 1% Penicillin-Streptomycin (Gibco, #15140122) was added. All imaging procedures were conducted under standard cell culture conditions (37 °C, 5% CO₂).

### Data preprocessing

The original image has a dimension of 10 × 2000 × 2000. We carried out maximum projection processing along the Z-axis. Subsequently, cell segmentation was accomplished via the mitochondrial channel utilizing Cellpose 3.0 software ^32,33^. Following this, cells whose centers of the region of interest (ROI) were located more than 70 pixels away from the image boundary were excluded. With the center of each retained ROI as a reference point, squares measuring 150 × 150 pixels were generated and employed to extract individual cells precisely. Across 280 compounds and 8-time points, over 400,000 images were ultimately selected for analysis from 282 drug treatment conditions (280 compounds and DMSO, NA). These images were then split into a training and test dataset at a ratio of 7:3.

### Structure of unsupervised clustering model

The neural network structure of the PB-scope is built based on a contrastive clustering framework ^15^. We divide the model into two branches, each subject to distinct data augmentation operations (Fig. 1c). Specifically, we elaborate on two data enhancement methods for each instance *X*_*i*_, through which we generate a set of 2N enhanced data samples, denoted as:

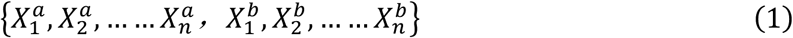

We utilize two independent MLPs to project features into the row and column space, where instance-level and cluster-level contrastive learning are conducted separately. We adopt ResNet34^49^ as the backbone network for feature extraction, which can effectively capture similar features of the samples with its powerful deep residual learning capability. In the training process, we employed the Adam optimizer^50^, with an initial learning rate of 0.0001. No weight decay or learning rate scheduler is applied. Due to time and memory limitations, the batch size is set to 128, and the model is trained for 200 epochs before testing (Supplementary Fig. 10).

### Multichannel color jitter

The data augmentation strategies ^23^ of PB-scope include resizing, cropping, converting to grayscale, horizontal flipping, Gaussian Blur, and Multi-channel Color Jitter, a data enhancement strategy for fluorescent images. In traditional contrast learning, the RGB channel intensities of natural images are usually correlated, as they often reflect the properties of the same object. However, when it comes to microscope images, each channel usually represents multiple cellular or tissue components tagged with distinguishable fluorescent markers^51^. Therefore, we incorporate an additional color perturbation model into the augmentation method to adjust the intensity of each channel differently to accommodate the noise introduced by individual laser source.

For an image *I* with dimensions *h* × *w* and color channels *c* ∈ {*R*, *G*, *B*}, the transformed image *I*′ is defined as:

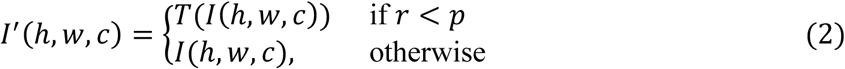

*T*(*x*) encompasses transformations of brightness, contrast, and saturation.

### Loss function of contrast clustering

The objective function for our model comprises two components: the instance-level and cluster-level contrastive losses as:

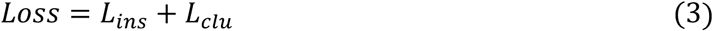

The instance loss *l_i_*and cluster loss 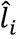 for a given sample *X_i_* is defined as:

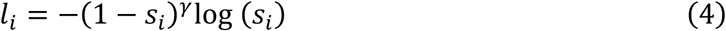

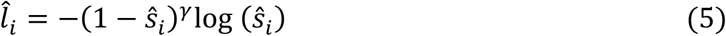

Where *s_i_* represents the similarity between sample pairs are defined as:

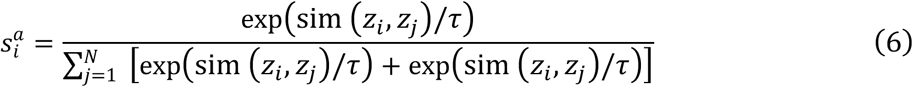

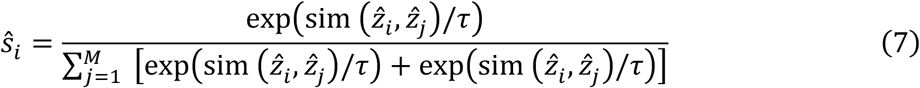

Wheresim (*z_i_*, *z_j_*) represents the cosine distance between the features of two different samples and *τ* serves as temperature parameters to regulate the degree of softness in each loss component. Overall, the instance-level and cluster-level contrastive losses are computed as follows:

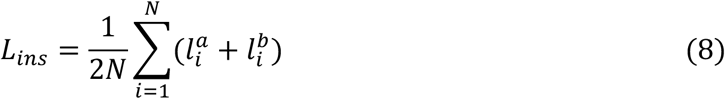

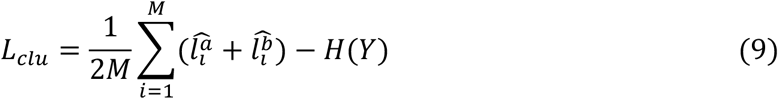

where *H*(*Y*) is the entropy of the cluster assignment probabilities.

### Clustering analysis

30 cell images were randomly selected for each drug from the test set to conduct clustering tests, and UMAP plots were generated. Forty-two drugs exhibiting noticeable clustering phenomena were identified and subsequently divided into five groups based on their UMAP distributions. Repeated experiments were carried out on two independent sets of images treated with drugs for 6 hours.

### P-body detection and quantification

For the detection of P-body, we generated a dataset by simulating cells and phase-separated particles (Supplementary Fig. 5). The cells were modeled as ellipsoids of varying sizes, and the background noise within them was assumed to follow a Gaussian distribution. For the phase-separated particles, their intensity is formulated as:

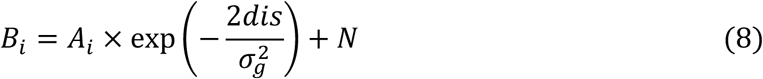

Where *B_i_* is the brightness, *Ai* is the parameter for the amplitude of the body’s intensity, and *N* represents the background noise.

We then took z-slices and generated corresponding labels for each. Subsequently, the generated dataset was used to train the YOLOv7 network. The images were first converted to 8-bit format. Subsequently, they were normalized to a range of 450 - 1000. Detection was performed to identify visible p-bodies, resulting in detection boxes. The confidence level for detection was set at 0.25. The number of p-bodies was then counted, and this count was divided by the number of cells to calculate the average number of p-bodies per cell. The protein expression level was determined by the average fluorescence intensity of the cell ROI subtracting the intensity of the background as:

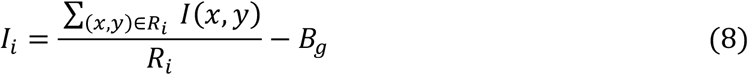

where *R_i_* represents the area of the cell ROI and *B*_3_ is the average intensity of background. These analyses were performed on three independent images, and their standard deviation was calculated.

### Immunofluorescence and microscope

HCT116 cells were seeded in 24-well plates covered with circle microscope cover glass (Nest, 801010) and cultured in DMEM with solvent or compounds. Then, cells were washed twice with cold 1xDPBS (Gibco, 14190144) and were fixed using 10% neutral buffered formalin (Bioss, C2034) at room temperature for 30 minutes, following permeabilization with methanol at −20 °C for 30 minutes, and blocking at room temperature for another 30 minutes using antibody dilution buffer (1xDPBS, 3% BSA). After the primary antibody was incubated overnight at 4 °C, cells were then incubated with the diluted secondary antibody for 1 hour at room temperature and 1xDAPI (MCE, HY-D0814) to stain nuclei. Imaging was performed on an inverted fluorescence microscope (BioTeK). As primary antibodies, we used DDX6 rabbit polyclonal antibody (Proteintech, 14632-1-AP) and EDC4 mouse monoclonal antibody (Santa Cruz Biotechnology, sc-376382).

### Quantification of P-body number in immunofluorescence images

Randomly selected immunofluorescence images from control, JAK1 siRNA, and JAK2 siRNA knockdown groups in HCT116 cells were analyzed to quantify P-body numbers. Individual cells with clearly defined boundaries and no overlap with neighboring cells were manually selected for analysis, while cells with ambiguous borders or overlapping regions were excluded. DDX6 channel was converted to 8-bit grayscale format. A consistent threshold range of 30 to 255 was applied across all images to detect and count P-bodies.

### RNAi knockdown

HCT116 cells were transfected with negative control (NC) or gene-specific siRNA oligos using Lipofectamine™ RNAiMAX Transfection Reagent (Cas No.13778150) and the cells were analyzed 60 hours after transfection. The siRNA sequences are CCGTATCTCTCCTCTTTGT for *JAK1* and GGAAATCTGAGGCAGATTA for *JAK2*.

### RNA extraction and expression analysis

The total RNA of cultured cells was extracted by RNA isolator Total RNA Extraction Reagent (Vazyme, R401) followed by isopropanol precipitation according to the standard protocols. Reverse transcription was performed with cDNA Synthesis Master Mix (Vazyme, R223). Quantitative real-time PCR was carried out using 2xSYBR Master Mix (Vazyme, Q711). Gene expression was normalized to *β-Actin* RNA. Primer sequences used for gene expression analysis are as following: *JAK1*_for 5’-GGTCAGCATTAACAAGCAGGACAA-3’, *JAK1*_rev 5’-AGCCATCTACCAGGGACACAAAG-3’; *JAK2*_for 5’-GGTGCTGAAGCTCCTCTTCT -3’, *JAK2*_rev 5’-CCGTGCACAAAATCATGCCG-3’; *β-actin*_for 5’-AGACTTCGAGCAGGAGATGG-3’, *β-Actin* _rev 5’-CAGGCAGCTCATAGCTCTTCT-3’.

### Statistics and reproducibility

The image-based drug screening was conducted in two replicates over 8 hours. The images from both experiments were combined for training and testing. During testing, 30 single-cell images were randomly selected from the test set for each drug to perform clustering analysis. No data were excluded from the study. The quantitative model was validated through three experimental trials, with the standard images verified by two experimental experts.

## Data availability

The single cell dataset utilized for model training, along with all model weights, is publicly accessible via Zenodo at https://doi.org/10.5281/zenodo.14591158. The simulation dataset utilized for P-body detection can be found via Zenodo at https://doi.org/10.5281/zenodo.15202103. The annotations of FDA-approved drugs were gathered from Drugbank. Further information and requests for raw data, scripts, and reagents should be directed to and will be fulfilled by the lead contact, T.T.

## Code Availability

The Pytorch codes for p-body drug screening can be found at https://github.com/todhc22skjicea/PB-scope. P-body simulations, detection, and quantitative analysis are available at https://github.com/todhc22skjicea/PB-scope-detection.

## Supplemental information

**Supplementary Table 1.** The compounds used in this study. The drug library comprises a collection of 278 FDA-approved kinase inhibitors, MG132, and thapsigargin.

**Supplementary Table 2.** Comparison of different unsupervised clustering algorithms. We selected 42 drugs from the original dataset that showed clustering characteristics in the embedding space and subsequently classified these drugs into five different groups (Group 1 to Group 5) based on their distribution patterns in the embedding space and constructed a new dataset named PB5 using the newly assigned labels for the five drug groups. This dataset was then utilized to evaluate the performance of the PB-scope. A variety of unsupervised clustering algorithms was selected as a comparison experiment. The experimental results show that PB-scope outperforms other common unsupervised clustering models in a clustering task involving fluorescent image data. The models were evaluated on the PB5 dataset; all deep-learning models were tested after 50 epochs of training.

| Method | ACC | ARI | NMI |
| --- | --- | --- | --- |
| Kmeans | 42.70 | 0.131 | 0.250 |
| AC | 46.81 | 0.157 | 0.275 |
| SC | 42.61 | 0.132 | 0.249 |
| NMF | 42.23 | 0.0198 | 0.079 |
| Simclr | 59.68 | 0.335 | 0.392 |
| CC | 61.35 | 0.408 | 0.455 |
| PB-scope(ours) | **63.27** | **0.430** | **0.479** |

**Supplementary Table 3.** Quantification of P-bodies in cells treated with compounds. Quantification of average P-body numbers and DDX6 expression levels per cell treated with different compounds. DDX6 expression levels were quantified by calculating the fluorescence intensity within the cells of three independent images.

**Supplementary Figure 1.**
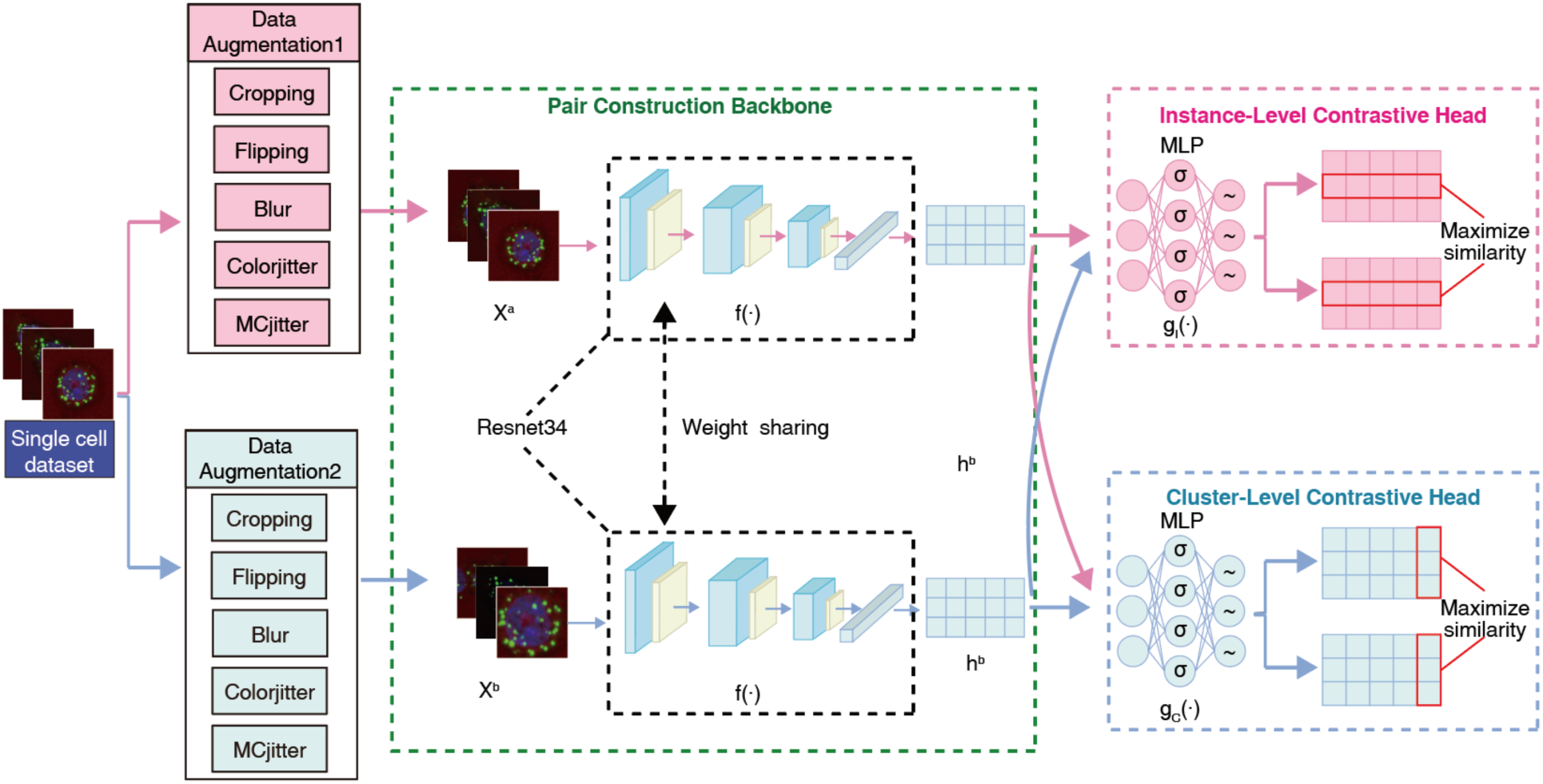
Contrastive clustering networks in PB-scope. The unsupervised clustering component of the PB-scope is implemented through a contrastive clustering framework. The model of contrastive clustering comprises two distinct branches, each subjected to different data augmentation operations. These augmentation methods include common methods such as resizing, cropping, converting to grayscale, horizontal flipping, and Gaussian blurring. Additionally, we integrate the MCjitter model into the augmentation approach to adjust the intensity of each channel differently, thereby accommodating the noise introduced by laser excitation across different channels. ResNet34 is selected as the backbone for feature extraction. Subsequently, it employs two independent Multi-Layer Perceptrons (MLPs) to map features into row and column spaces for independent instance-level and cluster-level contrastive learning (*σ* represents the ReLU activation function and *∼* represents the Softmax operation for the generation of soft labels.), aiming to capture the similarities between clusters and instances.

**Supplementary Figure 2.**
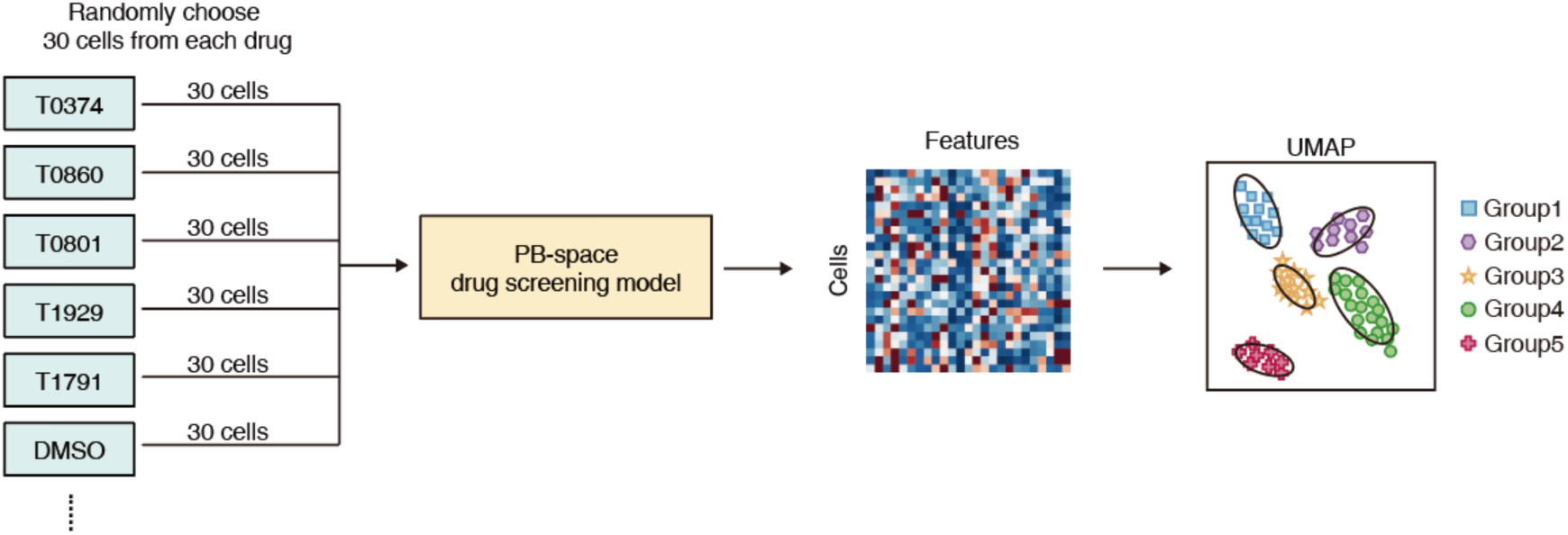
UMAP clustering methods. For cluster analysis, 30 cell images were randomly selected for each drug from the test dataset. Feature extraction was performed using the PB-scope to obtain the corresponding features for each cell. Subsequently, UMAP processing was applied to visualize these features, where each point on the UMAP represents a single-cell image with drug treatment.

**Supplementary Figure 3.**
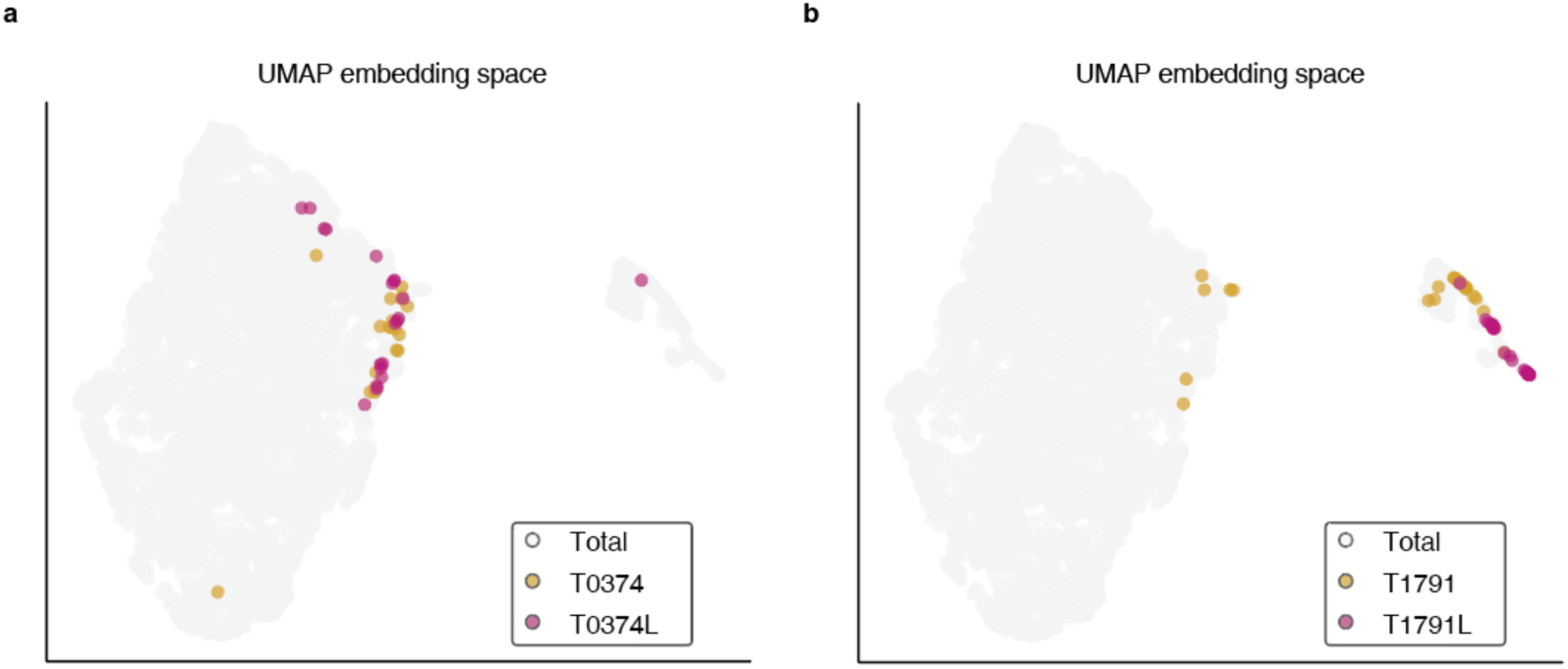
UMAP clustering analysis for structural analogs. UMAP cluster analysis shows that structural analogs (T0374/T0374L; T1791/T1791L) have close spatial proximity.

**Supplementary Figure 4.**
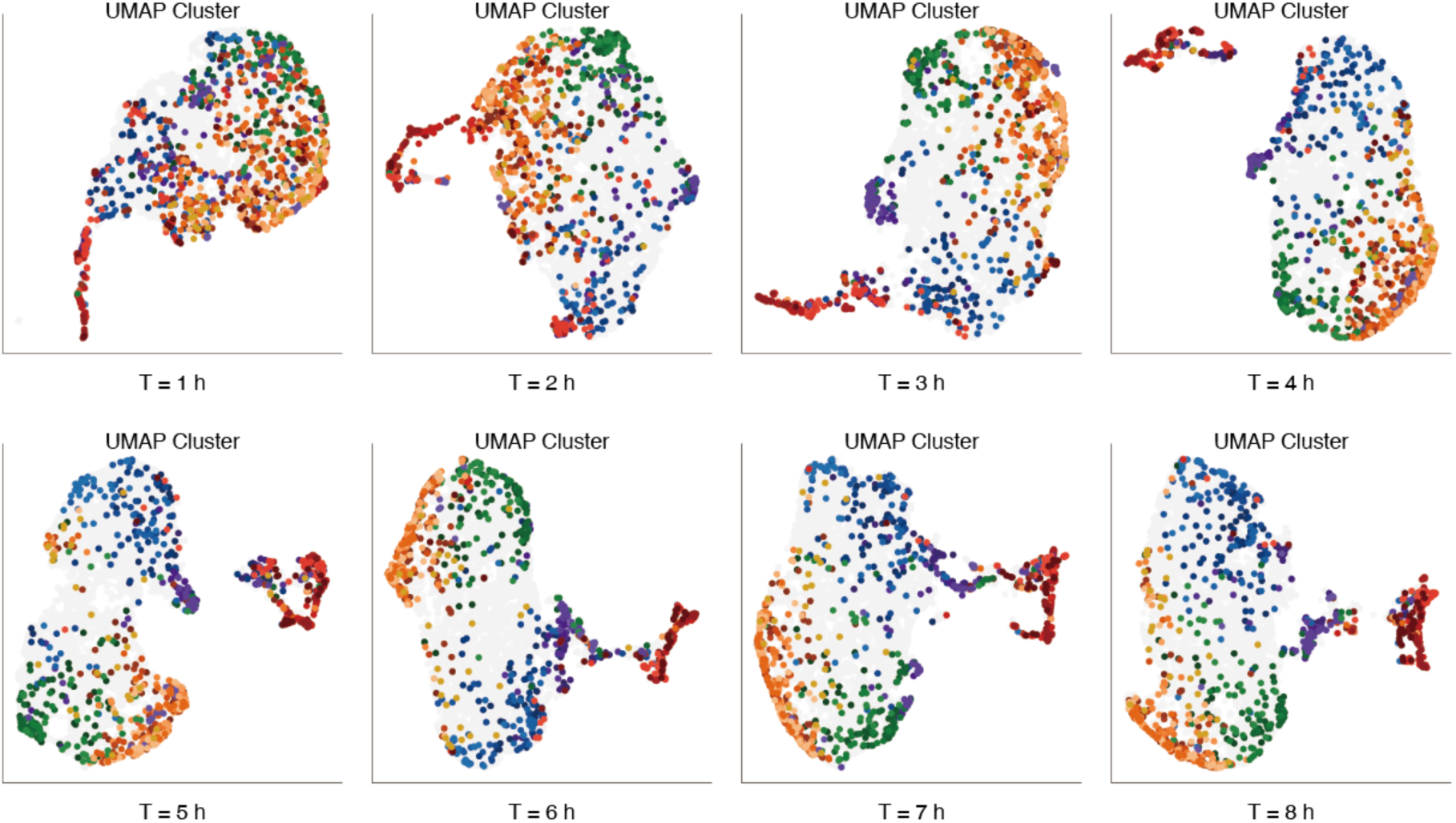
Time course data for UMAP clustering. To systematically investigate the effects of drugs on cellular phenotypic changes and the temporal evolution of structural features within the different groups, this study performed model training and cluster analysis on cellular phenotypic data collected at different time points (T = 1 – 8 hours). The experimental results show that the data points corresponding to the five groups of drugs transition from a uniformly overlapping state to a more cohesive state over time. This observation is consistent with the hypothesis originally proposed in this study, which is that prolonged drug exposure induces different phenotypic alterations that become more pronounced and differentiated over time.

**Supplementary Figure 5.**
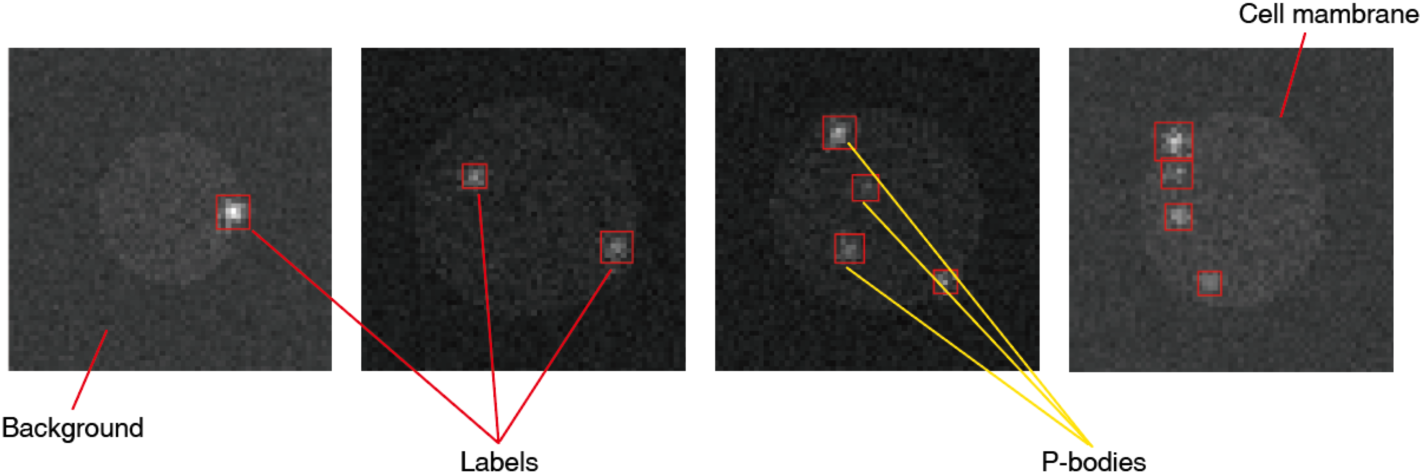
Examples of P-body simulation. The P-body particle simulation for generating datasets and corresponding labels encompasses various components, including the background, cell membrane, and intracellular particles. The fluorescence intensities of both the background and the cell are modeled to follow a normal distribution, ensuring a realistic representation of biological variability. Additionally, the intensity of the particles is designed to exhibit a Gaussian distribution related to their distance from the particle center, mimicking the natural spatial distribution of such particles within cells. For training and evaluation, the labels are stored in the YOLO format. This format allows for efficient annotation and retrieval of particle information, including their positions and class labels, thereby facilitating the application of P-body detection and localization.

**Supplementary Figure 6.**
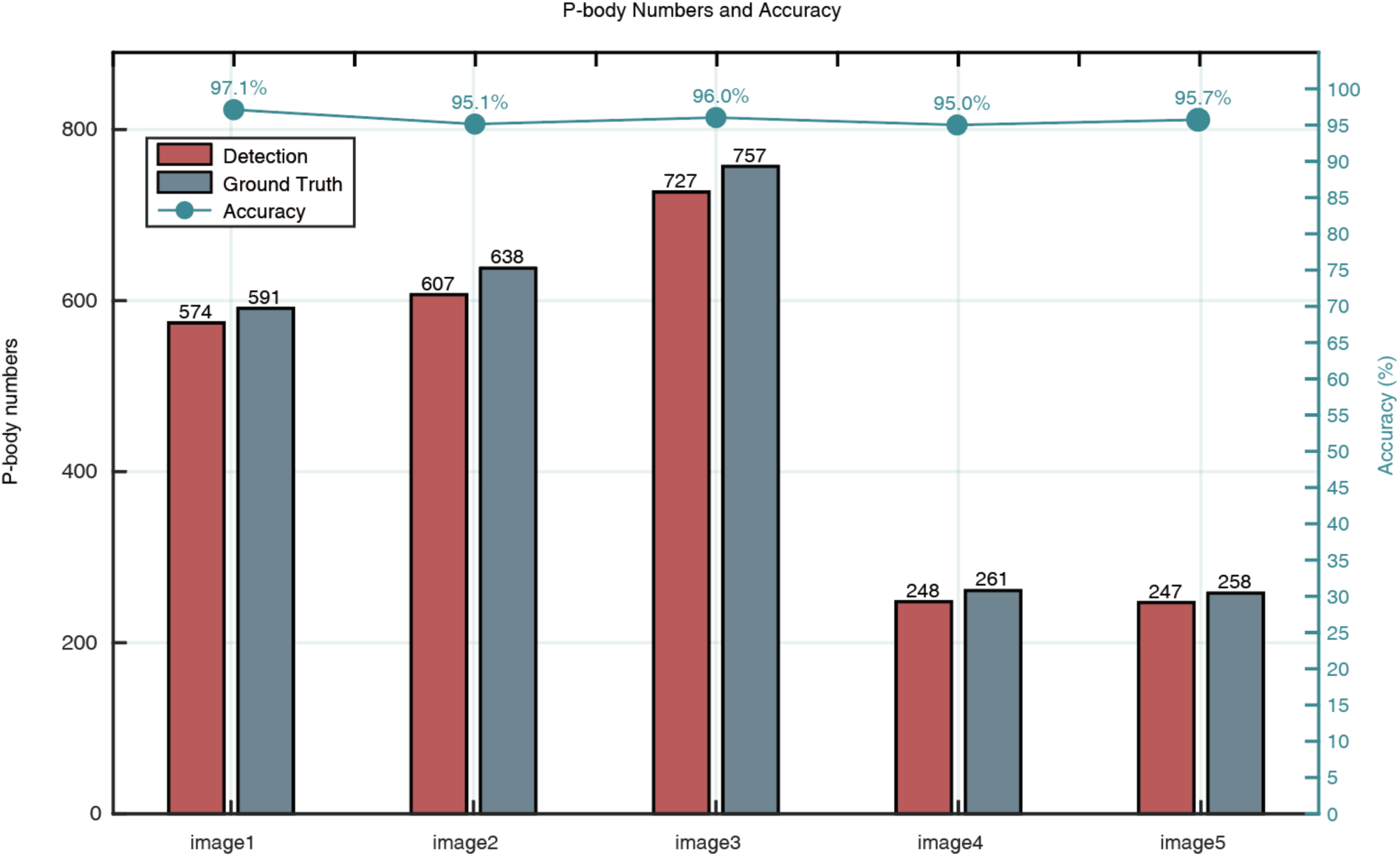
Validation for P-body detection model. To validate the accuracy of our simulated detection model, we selected five representative P-body images for comparison, which were manually annotated by two experts working in the related fields. Comparing the model’s detection results with the experts’ annotations showed that the model achieved an average accuracy of 95.92% in detecting P-body particles in colon cancer cells. This demonstrates that the model can reliably identify P-body and provides a powerful tool for P-body quantification.

**Supplementary Figure 7.**
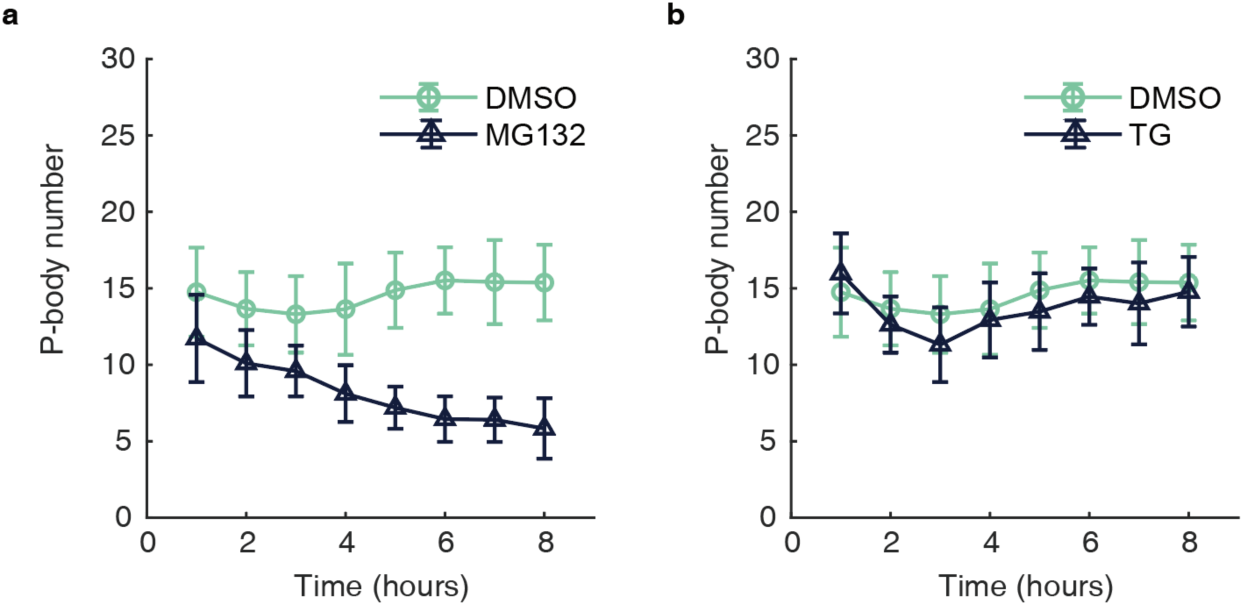
Quantification of P-body numbers for MG132 and thapsigargin treatment. MG132 induces a continuous reduction in P-bodies over 8 hours, while thapsigargin causes a temporary decrease in their number from 1 to 3 hours, followed by recovery after 4 hours. These findings are consistent with previous findings^39–41^.

**Supplementary Figure 8.**
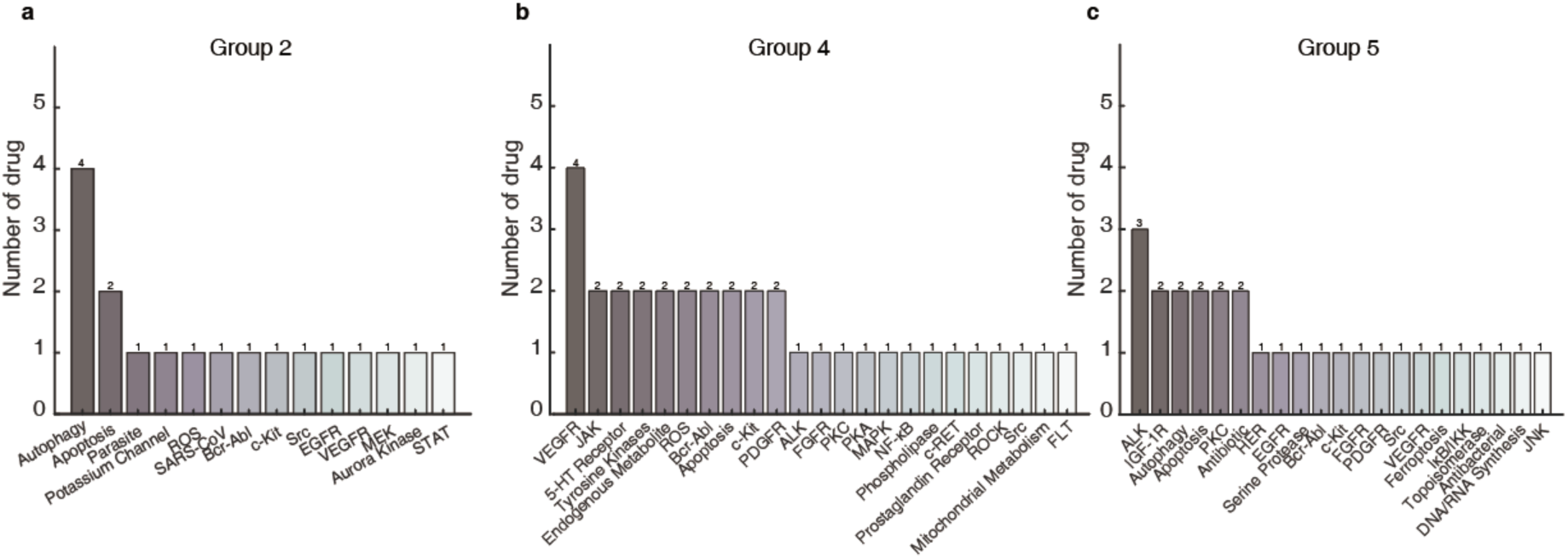
Mechanism of Action Profiling. Mechanism of Action (MOA) profiling for drugs in Groups 2, 4, and 5.

**Supplementary Figure 9.**
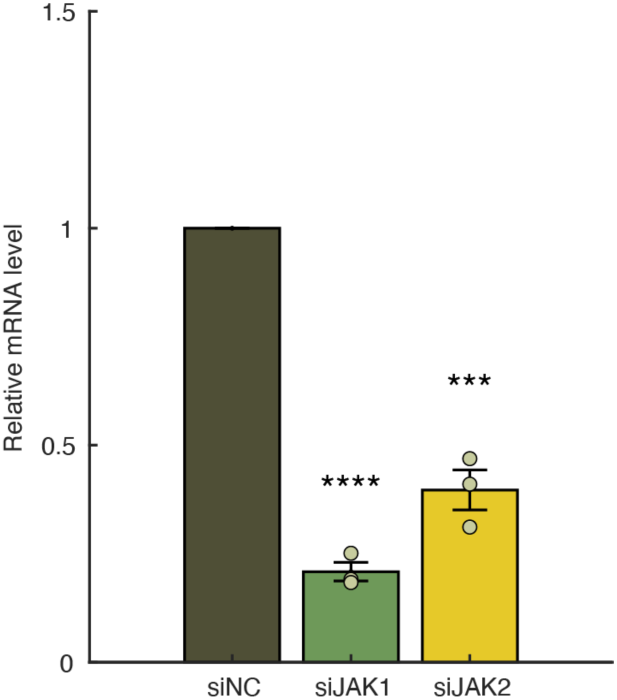
Confirmation of knockdown by RNAi. Knockdown efficiency is analyzed by qRT-PCR. The mRNA expression is depleted by the corresponding siRNA. Data (normalized by β-actin mRNA level) from three independent experiments are shown as mean ±SEM. The mRNA level in control cells is set to 1.0.

**Supplementary Figure 10.**
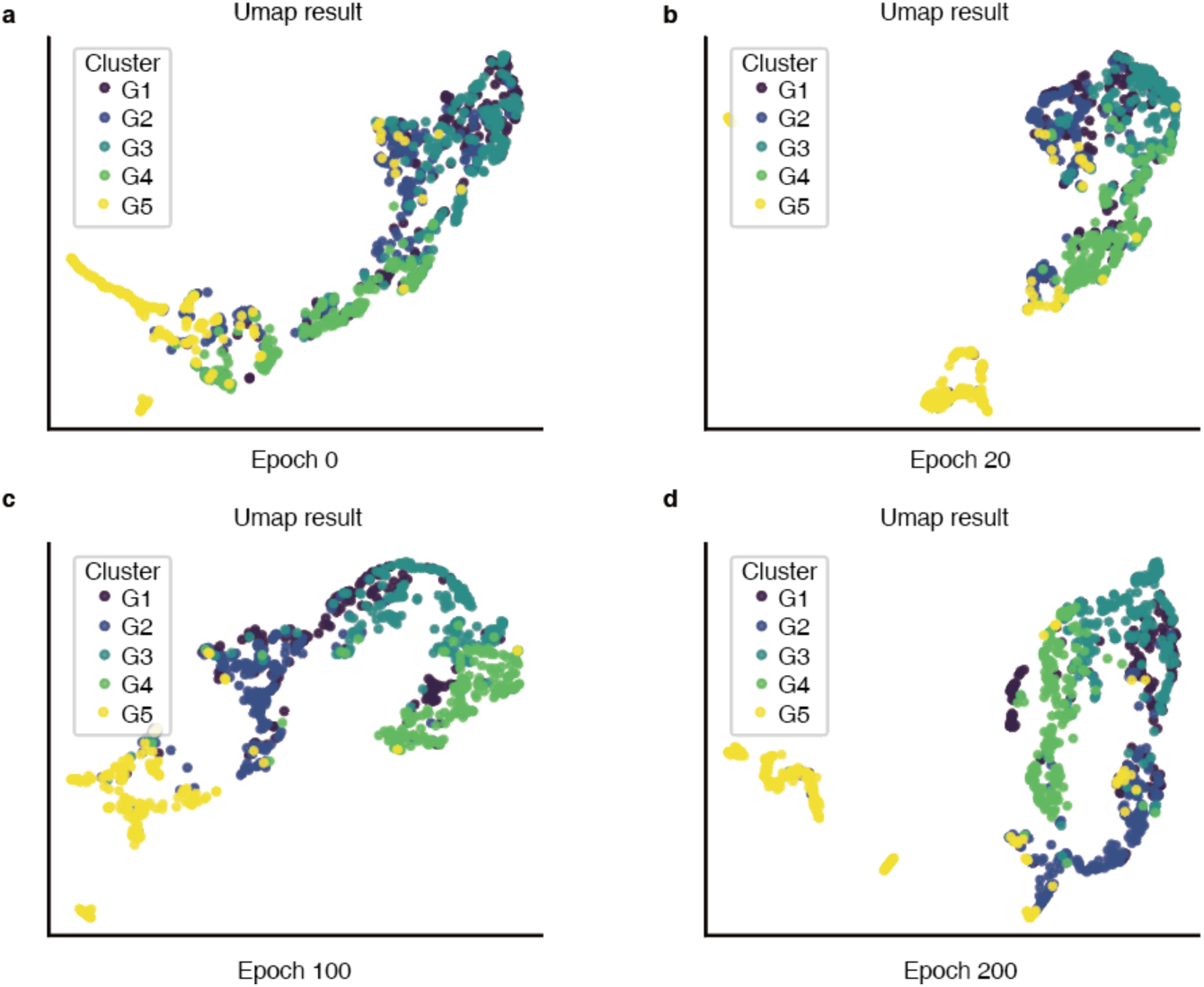
Training state of PB-scope. UMAP visualizations of the model’s feature space at four different time points throughout the training process. In these visualization analyses, different colors represent the true category labels of the phenotypic images (group 1 - group 5). At the initial stage of training, the distribution of cell image features in the UMAP space appeared haphazard. This suggests that the model’s ability to extract and distinguish drug-specific features is relatively weak in the early stage of the training process, making it difficult to effectively separate features from different drug classes. According to the model training, the distribution of features in the UMAP space becomes significantly more decentralized. Different clusters begin to form around each category, with increasing distances between the cluster centers. This phenomenon suggests that over time, the model gradually learns to extract and distinguish drug features. By the middle to late stages of training, the feature clustering stabilizes, and the location and shape of the cluster centers no longer change significantly. This stabilization indicates that the model has largely converged and reached a relatively stable level of proficiency in extracting and distinguishing drug features.

## Acknowledgments

We thank the members of Tsuboi and Chen laboratory for their helpful discussions and feedback on the paper. We thank X. Yang for annotating the P-body for ground truth data. This work was supported in part by the Key Research and Development Program of the Ministry of Science and Technology 2024YFE0102700, 2023YFA0914303, Shenzhen Science and Technology Innovation Commission WDZC20220811144737001, the Jilin Fuyuan Guan Food Group Co., Ltd, startup fund OD2021031C, Interdisciplinary Research and Innovation Fund JC2022008, Overseas Research Cooperation Fund HW2024009 from Tsinghua SIGS (to T.T.), National Natural Science Foundation of China 32470590, 32100613, Shenzhen Science and Technology Innovation Commission JCYJ20210324104605014, KQTD20180411143432337, Shenzhen Key Laboratory of Gene Regulation and Systems Biology ZDSYS20200811144002008 (to C.W.). We thank the authors of https://github.com/ManiadisG/DivClust^16^ for making their code public.

## Author contributions

Q.Z. W.C. and T.T. conceived and designed the project. Q.Z., X.P., D.P., M.Z., Y,L., Z.S., and L.F. performed wet experiments and analyzed image data. D.S. and Y.X. performed image quantification and computational analysis. D.S., Q.Z., Y.X., L.F., W.C., and T.T. wrote the manuscript.

